# Genome-wide Prediction of Potential Polycomb Response Elements and their Functions

**DOI:** 10.1101/516500

**Authors:** Morteza Khabiri, Peter L. Freddolino

## Abstract

The Polycomb-group proteins (PcG) and Trithorax-group proteins (TrxG) are two major epigenetic regulators important for proper differentiation during development (1, 2). In *Drosophila melanogaster* (*D. melanogaster*), Polycomb response elements (PREs) are short segments of DNA with a high density of binding sites for transcription factors (TFs) that recruit PcG and TrxG proteins to chromatin. Each PRE has a different number of binding sites for PcG and TrxG, and these binding sites have different topological organizations. It is thus difficult to find general rules to discover the locations of PREs over the entire genome. We have developed a framework to predict the locations and roles of potential PRE regions over the entire *D. melanogaster* genome using machine learning algorithms. Using a combination of motif-based and simple sequence-based features, we were able to train a random forest (RF) model with very high performance in predicting active PRE regions. This model could distinguish potential PRE regions from non-PRE regions (precision and recall ~0.92 upon cross-validation). In the process, the model suggests that previously unrecognized TFs might contribute to PcG/TrxG recruitment at the PRE locations, as the presence of binding sites for those factors is strongly informative of active PREs. A secondary regression model provides information on features that further differentiate PREs into functional subclasses. Our findings provide both new predictions of 7887 potential PREs in the *D. melanogaster* genome, and new mechanistic insight into the set of DNA-associated proteins that may contribute to PcG recruitment and/or activity.

## Author summary

During the development of multicellular organisms, the pattern of gene expression for every cell type must be established and then inherited through cell division. DNA *cis*-regulatory elements called Polycomb response elements (PREs) are major drivers of the spatio-temporal regulation of many genes during development. Recently, it has been reported that, depending on the cell type and condition, these elements could have dual functionality: a PRE can act as a silencing element in one cell type and an enhancer in others. Integrating all binding information of characterized TFs with DNA sequence features allowed us to construct a highly informative machine learning model to predict and classify PRE locations. Applying our model to *Drosophila* both allowed us to identify many new putative PREs (15-16 times the number currently known from experimental studies) and provides insight into the specific DNA-binding proteins that may additionally contribute to PRE function.

## Introduction

The cells of higher eukaryotes regulate gene expression at many different levels, from local transcriptional regulation to 3D chromosomal structure (3) and epigenetic silencing (4). The latter form of regulation is particularly important during development, as it allows gene regulatory programs to be maintained during cell division and differentiation throughout embryogenesis to form tissue, organ, and organism identity in higher eukaryotes (5–9). There are several major epigenetic regulators that contribute to maintaining a regulatory state over many cell divisions. Among them, Polycomb group (PcG) and Trithorax group (TrxG) proteins act epigenetically in functional opposition to maintain gene expression patterns as cells differentiate during embryogenesis: PcGs maintain gene silencing, whereas TrxGs act as anti-repressors to counteract PcG function (10–12). PcG proteins contribute to the formation of a specific type of repressive chromatin structure called a Polycomb domain which, for example, represses the expression of non-functional Hox genes during segmentation of the *Drosophila* embryo (13–15). Indeed, it has been shown that not only Hox genes, but also thousands of other genes essential for development, cell-fate determination, stem-cell pluripotency, and cell reprogramming are regulated by PcG proteins (14, 16). To date, three distinct *Drosophila* PcG protein complexes are known, including Polycomb repressor complexes 1 and 2 (PRC1 and PRC2) and pleiohomeotic repressive complex (PhoRC) (17, 18). Pho and *Drosophila* Sfmbt proteins (dSfmbt) form the PhoRC, which is the only Polycomb group protein complex that binds directly to DNA *cis*-regulatory elements called Polycomb response elements (PREs) in a sequence-specific manner (19–21). PRC2 complex is recruited by PhoRC to PRE regions. This complex has a methyltransferase enzyme that generates silencing H3K27me3 marks in a large domain, including regulatory elements like promoters and enhancers. Subsequently, the PRC1 complex recognizes the methylated regions and joins with PRC2 to form Polycomb complex at PRE regions (22, 23). Interaction of PRC2-PRC1-PhoRC complexes at PRE regions act as nucleation sites for Polycomb domains by formation of looping interactions (13, 15). PREs are typically thought of only as genomic elements that mediate PcG-dependent transcriptional silencing of associated target genes (17). However, recently published data from Erceg *et al.* (24) showed substantial additional richness of behavior, depending on sequence context and the exact developmental stage of the cell. Using PhoRC ChIP-seq peak coordinates, they categorized PhoRC peaks into four different classes: promoter, enhancer (developmental and chip-defined putative enhancers), intragenic (exon and introns of annotated genes), and intergenic (every other location). Further, they showed that, depending on the cell’s local environment, the PRE elements can have dual functionality during embryonic development -- the same PRE that recruits silencing PcG complexes in one type of tissue can act as an enhancer in different tissues.

Many PREs with similar functional properties are enriched with key motifs like Pho/Phol, Trl, combgap (Cg), Pipsqueak (psq), zeste (z), Dorsal switch protein (dsp), Grainyhead (grh) and Sp1/KLF motifs; however, the same elements show no preferred order, number, or spatial arrangement of consensus motifs in different PREs (11, 25, 26). Thus, there is no common rule to distinguish PRE from non-PRE regions due to the diversity in composition of PREs themselves.

Despite the difficulties involved, there have been several computational efforts to find a rule to predict PRE regions over the entire *Drosophila* genome (27–29). For example, Ringrose *et al.* considered 7 known motifs that exist in PRE regions and define a score for each possible pair of those motifs, then calculated the sum of the occurrence of every possible motif pair in each genomic window. Later, similar approaches were used to develop prediction algorithms by defining sequence feature properties, number of mismatch errors, and flexibility in motif pair combinations (27, 30). Although some of these predictions show high performance – e.g., a support vector machine (SVM)-based classifier had recall = 0.82 and precision = 0.96 (27) – the limited number of TF motifs and the small size of training data set (only 12 positive sites) relative to the overall diversity of known PREs makes it difficult to obtain a reliable computational prediction of PRE regions genome-wide. Therefore, recent ChIP-seq assays based on the presence of the PhoRC complex, as a main core of Polycomb group complex that recruits PRC1 and PRC2 to the PRE regions, provide a good opportunity to study more about PRE regions within the fruit fly genome.

To better computationally detect PREs, and to understand the relationship between PRE properties and their functions, we present here a novel computational approach to predict and classify potential PRE regions. Working under the hypothesis that binding of additional TFs in the vicinity of PRE-like sequences would specify Polycomb complex recruitment and activity, we built a database based on motif scanning that shows the potential binding of all possible DNA-associated proteins in each genomic region over the entire *Drosophila* genome. Using a RF model with the incorporation of the binding of all possible DNA-associated proteins for each potential PRE region as a main feature set, and combining it with sequence features, our approach was able to predict potential PRE regions with high consistency compared with ChIP-seq peaks and operationally known PRE regions. Previous computational studies showed that seven selected motifs corresponding to four TFs including GAF (two motifs), Pho (three motifs), engrailed-1, and z are informative in distinguishing potential PRE regions from non-PREs (27, 29). However, our study demonstrates that for prediction of potential PRE functions, more features/variables (motifs) are needed to improve computational performance and can provide useful information on PRE locations. We identify both known motifs for TFs such as Beaf-32 and stripe(sr), and newly inferred motifs specifically enriched in PhoRC binding sites, as being important features in identifying and classifying PREs. According to flybase annotations (31), most of these features correspond to proteins that are involved in key biological processes such as development, cell organization, and proliferation. Our results suggest that the formation of functional PREs in a given tissue type is driven in part by contributions from several accessory DNA-binding proteins active in that particular tissue type; thus, we have simultaneously obtained a highly effective specific model for finding PREs in our condition of focus (mesoderm cells and in 4-6h and 6-8h windows of embryogenesis), and also obtained important insight into strategies that will enable prediction of active PRE locations under other conditions in which the set of active accessory DNA-associated proteins may be different.

## Results

We developed a new machine-learning framework based on the RF algorithm to predict potential PRE regulatory elements and their functions within a regulatory network over the entire *D. melanogaster* genome (Fig 1). As a key starting point, we constructed a database to classify the strength of binding of known TFs and other DNA-associated proteins to each specific location of the *D. melanogaster* genome based on combined information from all available sequence motifs in the modENCODE (32, 33) and CisBP databases (34). Subsequently, based on a recent ChIP-seq assay for nearby Pho and dSfmbt binding sites to obtain a reliable training set for active PREs, we developed models to predict PRE locations and to categorize them into different classes.

**Figure 1.**
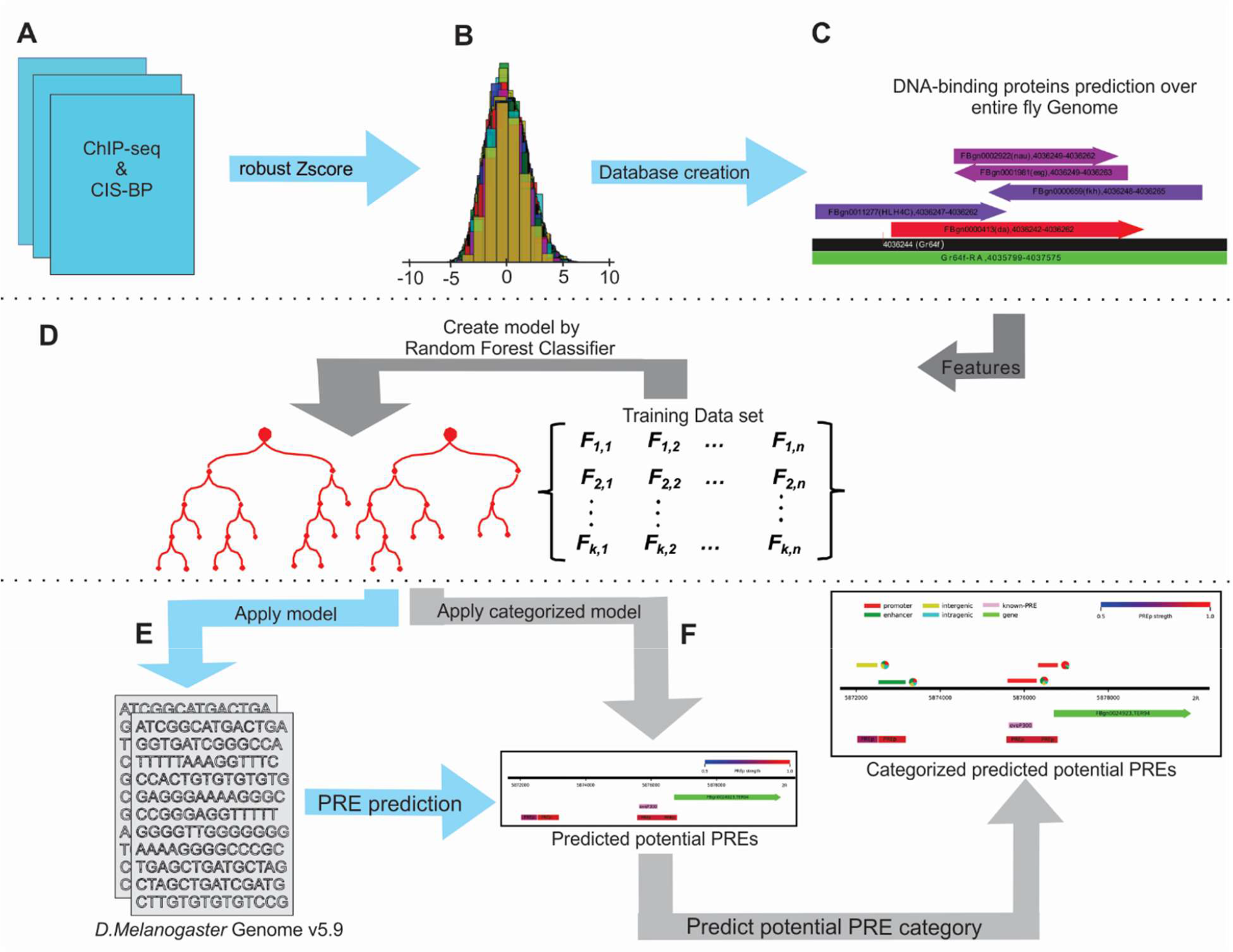
Flowchart of procedure for prediction of PREs. (a) Collect all known DNA-binding protein motifs in *D. melanogaster*. (b) Establish background distributions for each motif by checking for binding across the entire genome and calculate the robust zscore (rz-score) for each motif at each site. Each histogram shows the population of rz-scores for one DNA-binding protein. (C) Prediction of binding sites for each DNA-associated protein based on rz-score. The black line indicates genomic locations. The name of the gene overlapped with that particular of the genome is shown in parentheses. The green line shows the gene region in that particular location of the genome. The arrows above the genomic location (black line) indicate predicted binding sites, with the strength (based on rz-score) of binding sites indicates by color intensity from blue (as weakest) to red (as strongest) binding. The arrows indicate strand direction relative to the reference motif (+, to the right) and (−, to the left) (D) Using rz-scores and sequence features, construct two random forest models to predict and classify potential PREs. (E) Apply RF model over entire *D. melanogaster* genome to predict potential PRE regions. (F) Apply classified model on predicted potential PREs in order to determine the chance of each category (i.e promoter, enhancer, intergenic and intragenic) at each potential PRE regions.

### Constructing predictive database for known TF binding motifs

Because DNA-binding proteins such as TFs apply their functional roles by binding to regulatory elements such as promoters, enhancers, insulators, PREs, and TREs, we expected that the combination of DNA-associated protein information along with sequence properties would be optimal for distinguishing PRE regions from the rest of the *D. melanogaster* genome. We already know that each PRE has a different set of DNA-binding proteins with different order and arrangement based on operationally known PRE regions (25). Thus, we hypothesized that using the combination of all possible DNA associated proteins bound to each potential PRE region and considering sequence features such as nucleotide composition could allow us to construct a model to identify PREs throughout the genome. To test this hypothesis, we collected all available DNA-associated protein position weight matrices (PWMs) available in the CIS-BP database (34). We also gathered binding data from ChIP-seq assays in both modENCODE(32, 33) and literature sources (35) to construct the PWMs for DNA-associated proteins which do not exist in standardized databases (S1 Table and S2 Fig). To extend our list with motifs directly relevant for PRE regions, we applied *de novo* motif searching on the 994 ChIP-seq peaks found in (24), as described in the Methods section. The discovered motifs, which represent sequence features enriched in the PhoRC peaks, are listed in S2 Fig, along with predictions of what protein corresponds to each motif using Tomtom (36). Since the peaks from which new motifs were inferred belong to PhoRC complex binding regions, we named the newly identified motifs in these regions Inferred Pho Related Motif 1 (IPRM1), IPRM2, IPRM3, etc.; note that this naming does not mean that a particular motif corresponds to binding of Pho or a similar protein.

To identify potential binding sites for each DNA binding protein in our database, we calculated all possible binding scores of each PWM (x_i_) for binding at each possible location across the *D. melanogaster* genome. Thus, for each DNA-associated protein we would have a unique distribution of potential binding scores X_j_ (Fig 1B). Then the robust zscore (rz-score) at each position is calculated for each DNA-associated protein in our collection separately. The rz-score is calculated at each position by reference to the median and median absolute deviation (MAD) of the distribution of protein binding scores for that protein across the entire genome. The rz-score is calculated by the following formula:

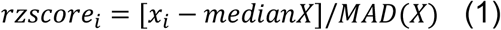

Therefore, we constructed an exhaustive database that predicts the strength of a motif match at each *D. melanogaster* genomic location relative to the overall distribution of scores for that motif (Fig 1C and S3 tables). Given the extensive coverage of TFs, we expected that these databases would cover a significant number of cis-regulatory sequences as well as components of PREs to regulate gene expression. The use of motifs, rather than primary ChIP-seq peaks, to inform our model allows us to incorporate information from methodologies such as SELEX methods, and avoids bias toward only sites bound under the particular conditions in which ChIP datasets were gathered (which do not always overlap with our condition of interest).

### Predictive Model for Active PRE Locations

Our goal was to model the identities of PRE regulatory elements and their roles, distinguishing PREs from non-PRE sites throughout the genome. The RF (37) algorithm was used for the classification because of its robustness in the face of potentially irrelevant features. We developed two RF models based on the rz-score database: one model to distinguish potential PRE from non-PRE regions (Fig 1E), and a second model to predict the class of each potential predicted PRE as being a promoter, enhancer, intergenic, and intragenic (Fig 1F). The motivations and goals of the two models differ somewhat; the first is intended primarily for *de novo* PRE discovery, whereas the second is intended mainly for regression purposes to identify features that distinguish the different PRE subtypes, since their sub-classifications can also be inferred by position alone(24). Each model had two major feature components: (i) rz-scores for those motifs existing at each potential PRE region, and (ii) DNA sequence features.

### Construction of the training set

To train an RF model for identification and classification of PREs, we established a training set consisting of two parts: 1) A positive dataset for PRE sequences, and 2) a negative dataset for non-PRE sequences. For the positive training dataset, we use a recent ChIP-seq data set (24) for the overlapped Pho and dSfmbt binding sites as a central PRE-binding complex; we thus develop our model under the assumption that the presence of PhoRC binding is indicative of an active/functional PRE. The size of the potential PRE regions (*i.e.* peak ranges) varied from 100 to ~1000 bps with the highest frequency at ~150 bps. Thus, we selected the summit of each peak as a midpoint and extended it 150 bp to the left and right to generate a positive region. As a negative data set for training our classifier, an equal number of potential PRE sequences with the same length were randomly selected from whole *D. melanogaster* genome. Taken together, we constructed a sequence collection containing 994 known potential PRE sequences (as the positive training set) and 994 non-PRE sequences (as the negative training set). To test the performance of model the data was split to 75% of the data for model generation and selection, and 25% of data for testing of the final model – the latter set was never used for training the model. As described below, we also show that the precise identities of the negative set sequences did not strongly affect the model.

### Construction of a random forest model to identify potential PRE locations

Our RF model for identifying PRE regions was implemented as described above and in the methods section. We used two measures to evaluate the performance of each model. The first is the area under the receiver operating characteristic curve (AUC-ROC), where a perfect model would have AUC-ROC = 1, and the second one is the area under the precision-recall (P-R) curve, where precision is the proportion of correctly predicted potential PREs and recall is the proportion of potential PREs that are correctly predicted. We construct our final model based on tuning RF parameters. After tuning RF parameters, our fitted models for identifying PREs from non-PREs showed AUC=0.92 for both ROC and P-R curves, taking averages over 5-fold cross-validation (Fig 2, GroupA; see below for a discussion of the different feature Groups tested).

**Figure 2.**
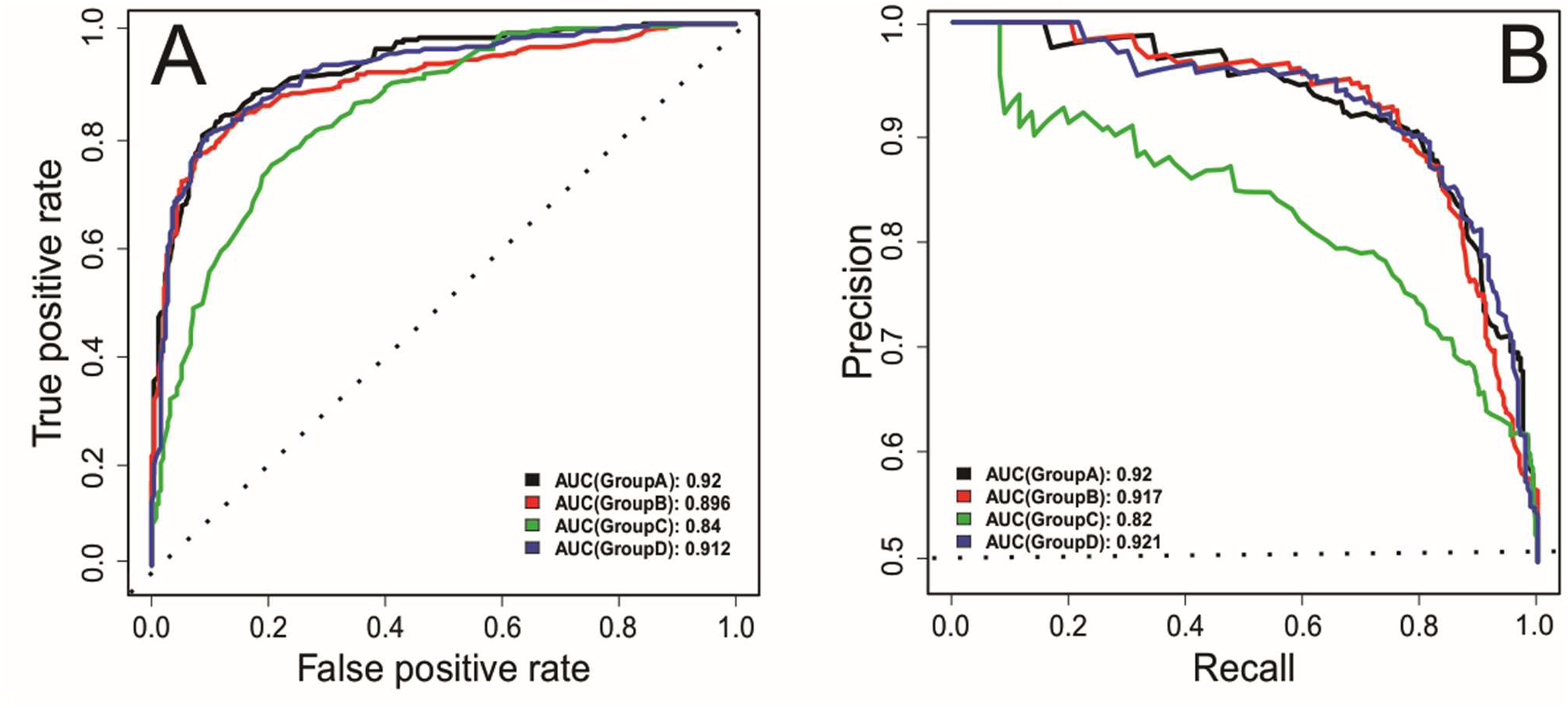
Evaluating performance using receiver operating characteristic (ROC) and precision-recall (PR) curves on the (withheld) testing data set. (A) ROC curves for models predicting potential PREs using all motifs available in our predicted DNA-associated proteins binding database (Group A); Pho, Pho related motifs, Trl and Trl related motifs (Group B), all motifs in Group A except motifs in Group B (Group C); or the motifs in ChIP-seq assay as listed in Fig. S2 (Group D). AUC indicates the area under each curve. The groups are defined in more detail in the text. (B) As in panel A, plotting P-R curves. Dashed lines indicate a hypothetical null model’s performance.

Looking at DNA-associated protein sites within our positive training data set, we find that 308 out of 378 DNA associated proteins in our predicted database exist in at least one PRE (S4 Table). Among the motifs present for each DNA-associated protein at each PhoRC binding region, the maximum rz-score among all motifs of a protein present at each sequence with rz-score greater than or equal to 3 were retained and considered as significant in binding to the region.

In addition to Pho and dSfmbt, there are several major motifs associated with PRE/TRE functions that are already known. These include motifs for the sequence-specific DNA binding proteins Pho (38), Trl (39), EN1 (40) and z (41). However, the genome wide ChIP-seq assays for these TFs reveal that these factors also bind in many locations in the genome outside of PREs and are involved in other regulatory networks. Thus, there must be additional features and motif binding contributing to the establishment of PRE/TRE regions. To discover if any other DNA-associated proteins other than Pho, Trl, dSfmbt, EN1, and z are involved in specifying potential PRE regions, we divided our feature set into four categories. (1) Group A includes max rz-scores of motifs for all 308 DNA-associated proteins which appear in ChIP-seq peaks of PhoRC complex (24), as well as DNA sequence properties. (2) Group B ONLY includes max rz-score of known Pho, Pho related motifs, Trl and Trl related motifs, and sequence properties. This includes Pho and Trl motifs provided by CIS-BP database, and PhoRC motifs IPRM5, IPRM6, IPRM7, IPRM14, IPRM17, IPRM23, and IPRM25 (S2 Fig). (3) Group C includes everything in Group A except for the motifs in Group B; thus, this group represents non-Polycomb/Trithorax related motifs which might, in a naïve understanding of PREs, be expected to yield no information. Finally, (4) Group D includes only the motifs found by motif analysis of PhoRC ChIP-seq data, i.e. all motifs described in (S2 Fig), and sequence properties. We fit an RF model for each individual group separately and then calculate the model performance on our withheld testing set based on ROC and P-R curves (Fig 2). Based on the AUC values in both ROC and P-R curves, the performances of models A, B and D are very similar. It is striking that model C, which excludes all important known motifs corresponding to Pho, Trl, and z, still shows high performance in distinguishing potential PRE regions from the rest of the genome, demonstrating that the presence of binding sites for several accessory TFs or other DNA associated proteins must also be informative of PRE status (albeit not as much so as the motifs for known core elements of PREs).

We applied feature importance analysis to identify which motifs from our database were most informative for identifying PREs from non-PRE regions. We considered two relevant measurements: mean decrease in accuracy (MDA) and mean decrease in the Gini index (MDG). A higher value of the MDA or MDG score indicates higher importance of the feature to our model. Fig 3 shows the MDA and MDG scores for the top 30 important features involved in constructing models based on Group A features. Aside from Trl, Pho, and their related motifs, several additional TFs such as Beaf-32, CTCF, Dref, IPRM11 (disco), IPRM9 (Zif/Zipic) and sr appear as important features to create the best possible model. Except for Beaf-32 and CTCF, which are very important in chromatin remodeling (42, 43), the rest of the newly implicated factors are involved in *D. melanogaster* development and the embryogenesis process (31). The important features involved in constructing models based on Groups B-D are depicted in (S5 Fig). Comparing important features highlighted prominent roles of similar sets of DNA-associated proteins within potential PRE regions, regardless of the exact feature set chosen.

**Figure 3.**
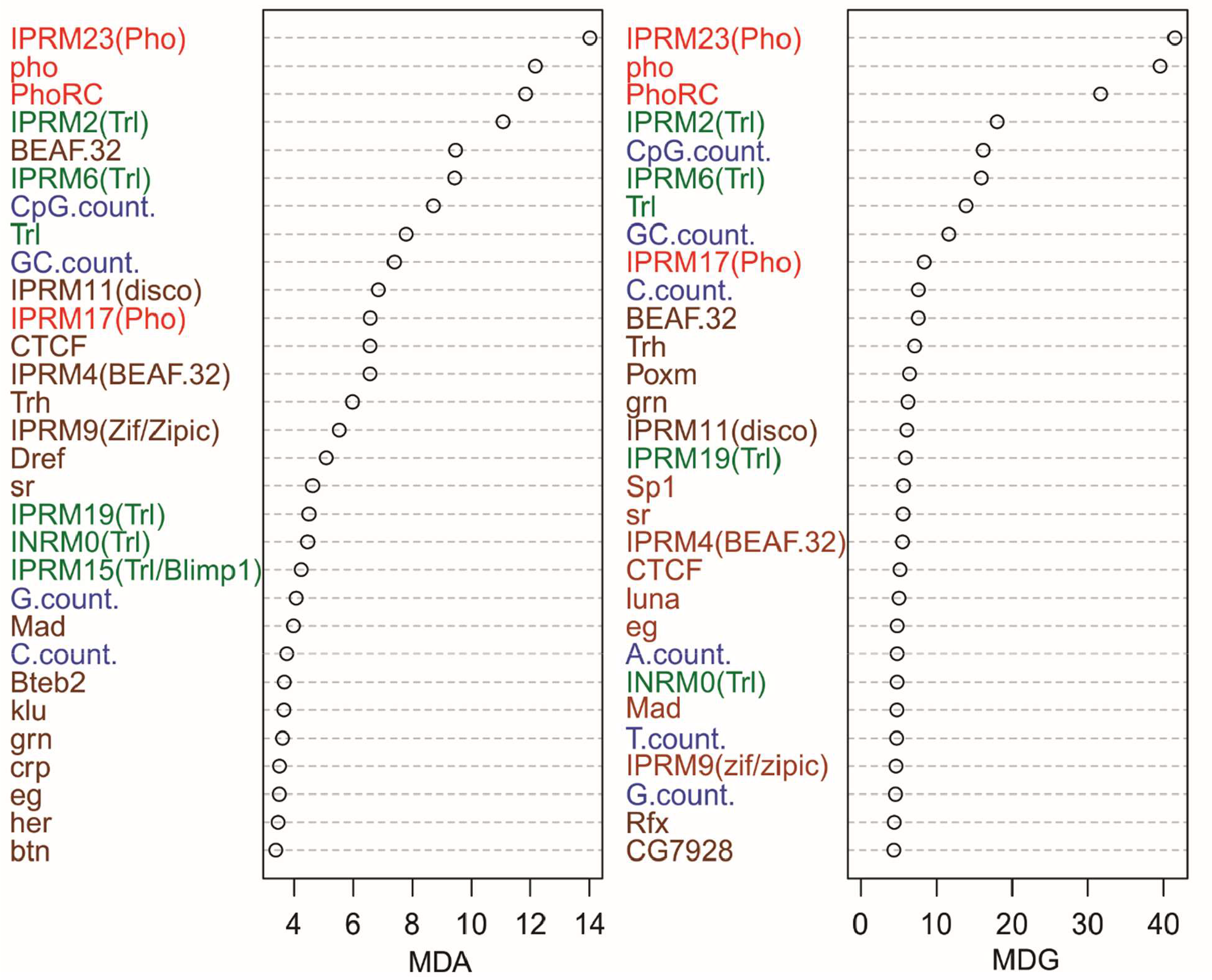
Variable importance table for the RF model based on all motifs (that is, Group A). Mean decrease in accuracy (MDA) and mean decrease in Gini coefficient (MDG) of effective factors in RF model are shown. Trl and Trl related motifs are shown in green, sequence features in blue, Pho and Pho related motifs in red, and other features (predominantly other motifs) in brown.

In order to identify which DNA-associated proteins occurred more frequently within PRE/TRE regions, we show the distribution of rz-scores of the 15 most important variables in both PRE and non-PRE regions (Fig 4). Testing for higher rz-scores in the positive dataset when compared with the negative set was done using a one-sided Wilcoxon test; we then present q-values based on false discovery rate (FDR) adjusted p values via the Benjamini-Hochberg method(44). The results of our significance tests reveal that all of these features occur significantly more often in the positive data set (i.e in PRE regions). Thus, one might expect that PRE regions could be identified by searching for these DNA-associated binding sites. To ensure that the model based on Group A was not biased by the randomly selected negative dataset, we performed random selection of the negative dataset from the whole genome a total of ten times, thus creating 10 different individually cross-validated models that all have the same positive dataset but different in negative datasets (S6 Fig, panels A and A’), selecting and withholding a random quarter of each set as a test set (*n.b*. the same positive examples were withheld in each case). Comparing the AUC of P-R and ROC curves show that all constructed models have similar accuracy and performance. The average of these curves is very close to our tuned RF model which is a strong support of the performance of our model. In addition, we applied our main model (without retraining) to the test dataset associated with each of those other training data sets. The positive examples of each test set are the same, and neither this dataset nor the negative datasets of the 10 different test sets are included in the training dataset of our main model. The B and B’ panels in Fig S6 indicate the performance of the main model on these 10 different test datasets. The similar AUC values under both ROC and P-R curves provides further evidence for the robustness of our main model, and the lack of any bias due to the selection of the negative training data.

**Figure 4.**
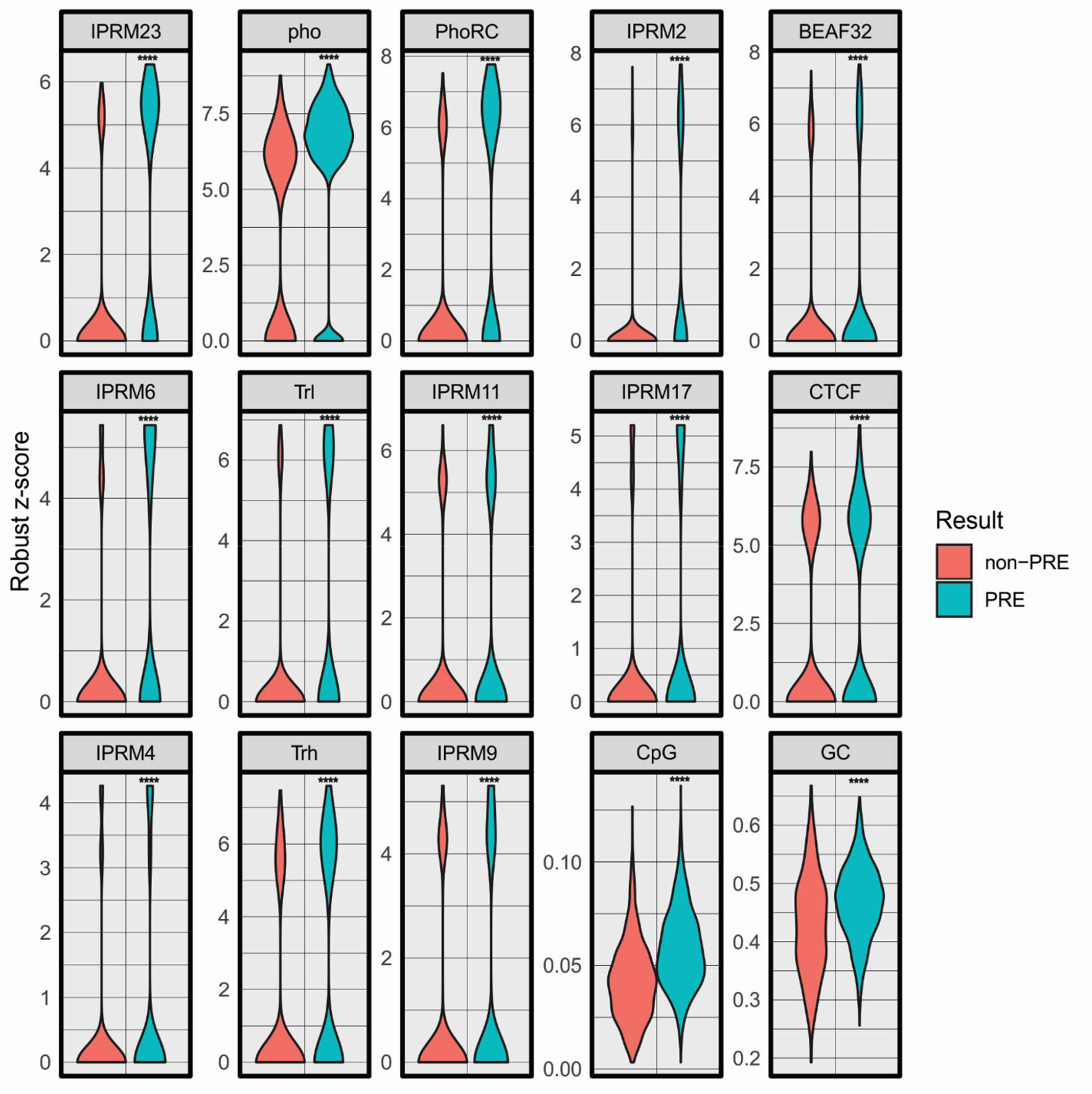
Distribution of rz-score of top 15 important variables, selected from Figure 3, in PhoRC and non-PhoRC biding regions. For each feature we performed a one-sided Wilcoxon rank sum test with an alternative hypothesis that the distribution for the PRE regions is stochastically greater than that of the non-PRE regions. The stars show the levels of significance using q-values (FDR-corrected p-values). If a q-value is less than 0.05 it is one star (*), less than 0.01 it is two stars (**), less than 0.001 it is three stars (***) and less than 0.0001 it is four stars (****).

### Genome-wide application of potential PRE location predictions

The high performance of our model on different test sets (with the same positive and different negative data sets, as mentioned above) encouraged us to use it to search for PRE/TREs throughout the *Drosophila* genome. We applied our model to 300 bp windows moved in 50 bp step intervals across the entire sequenced *D. melanogaster* genome v5.9 obtained from flybase(31). In each window, the possible binding motifs and sequence features were calculated. In each window the maximum rz-score were selected if more than one motif hit was found for a given DNA-associated protein. Then, our main RF model (described above) was applied on each window. The output at each location is the fraction of trees in the RF model voting that a given site represents a PRE, which can be taken as a confidence score for the prediction that the region indeed represents a PRE.

### Comparison with Additional ChIP-based PRE Datasets

To provide an unbiased evaluation of our genome-wide model and identify reliable confidence score thresholds for the identification of new PREs, we take advantage of two experimentally identified gene lists (8, 45) that are regulated by PRE/TREs activity, but were not directly incorporated into our model training. Schuettengruber et al. (45) measured binding of Pc, ph and H3K27me3 member of Polycomb group proteins in the *D. melanogaster* embryos within 4-12h of embryogenesis window, while Schwartz et al. (8) measured Pc, E(z), and Psc as PRE markers to determine PRE elements in S2 cultured cell line. For each data set, genes with overlapping transcription start sites (TSSs) with these regions were selected as target genes that are regulated by PRE elements. The gene names in both lists (obtained from the primary publications noted above) were adjusted to the gene name of genome version 5.9 obtained from flybase(31). To determine a best confidence score cut-off for our model, we defined a range of thresholds between 0 and 1 with intervals of 0.01. At each point, we did a binary classification to identify potential PREs throughout the genome: the sites with voted scores greater than that threshold would be considered as potential PREs, and the sites with scores less than that threshold would be considered non-PREs. If several windows adjacent to each other were predicted as potential PRE sites, they were all combined into a longer potential PRE. Then, we defined genes that have one or more potential PRE regions within 5000 bps upstream of their TSS as genes that are regulated by PRE function. For determining the optimal threshold, we used the Matthews correlation coefficient (MCC), which measures the balance of prediction sensitivity and specificity (46), to compare our model predictions with the gene sets identified in the Schuettengruber and Schwartz papers. The MCC is a correlation coefficient between the observed and predicted binary classifications, and it returns a value between −1 and 1. A coefficient of 1 indicates a perfect prediction, 0 an average random prediction, and −1 a perfect inverse prediction. We note that neither of the experimental data sets used here are expected to correspond perfectly to the active PREs in our training set – they may be sensitive to protein binding events that do not actually indicate active PREs, are subject to loss of resolution due to only providing gene-level identification of targets, and are taken from different tissues under different conditions – but at least some overlap of targets is still expected. Thus, we calculate the MCC coefficient at each point within the range of threshold between 0 and 1 with 0.01 intervals (S7 Fig). According to MCC plot, the threshold ~ 0.9 has the highest MCC score while the threshold 0.8 also gives reasonable performance (and substantially higher recall); on the training data described above, the 0.9 threshold gives a recall of 0.03 and precision of 1.00, whereas the 0.8 threshold gives a recall of 0.26 and precision of 0.98. These scores provide optimal overlap between our PRE predictions and those inferred from the two experimental data sets considered here, neither of which was used directly in training our model. Thus, in most of our analyses below, we report data on our genome-wide PRE predictions at confidence score thresholds of 0.8 and 0.9 to represent low-confidence and high-confidence predictions.

Using thresholds of 0.8 and 0.9, we identified 7887 and 563 locations as potential PRE regions that cover approximately 3.93 and 0.34 % of the whole *D. melanogaster* genome, respectively (S8 Table). Calculating the overlap of the positions of predicted PREs with the 5000 bp upstream of the genes reveal that there are 378 genes whose expressions are potentially being regulated by PRE elements in *cis* (S9 Table); an unknown additional number could be affected via enhancer activity. Fig 5 shows the overlap of our predicted genes and the two experimental datasets described above, demonstrating strong enrichments in overlap between our predictions and both experimental data sets. Since our potential PRE prediction is based on specific cell type (mesoderm cells) from embryos in 4-6 and 6-8 h of embryogenesis, perfect overlap with the two experimental studies should not be expected. The statistical significance of the overlap of each experimental dataset with our predicted gene list at both 0.8 and 0.9 cutoffs was estimated utilizing a permutation test. Laying at the center of all gene sets under consideration, 34 genes and 89 genes are in common between our predictions and both validation data sets using our 0.8 and 0.9 cutoffs, respectively (S9 Table). According to the gene annotations in the Flybase database(31), each of these genes is involved in biological pathways important in the developmental process, which is consistent with the expected roles of PREs in regulating key developmental processes of an organism. Thus, our model provides strong performance in genome-wide PRE predictions that are consistent with experimentally inferred sites while providing predictions of hundreds of additional active PREs that are likely functional in embryos 4-6 h and 6-8 h after fertilization.

**Figure 5.**
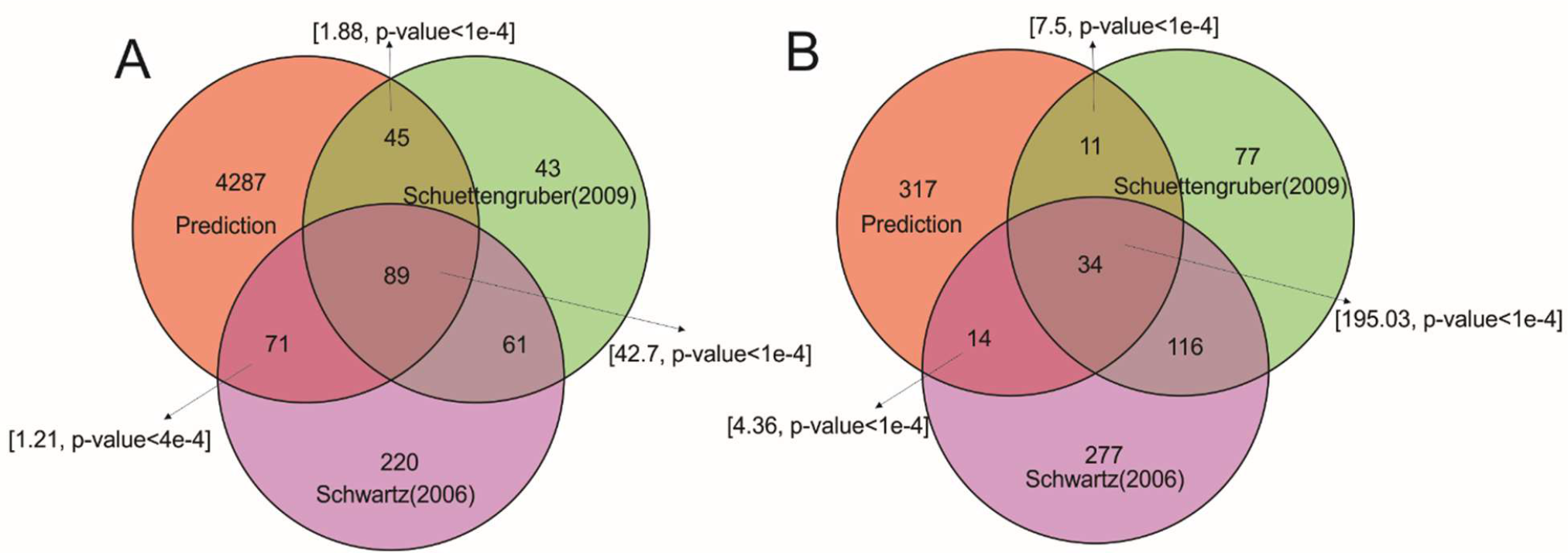
Venn diagram of overlaps between different lists of PRE-regulated genes at cutoffs 0.8 (A) and 0.9 (B). The Venn diagram of the target genes by potential PREs with two experimentally known gene list predicted through ChIP-on-chip technique by(45) and (8) as described in the text. Overlaps are based on the cutoff 0.8 and 0.9 for our RF model. The arrows and brackets indicate the gene enrichment (relative to expectations assuming no correlation) and corresponding p-value in each intersection (calculated using a permutation test as described in Methods).

### Function prediction of potential PREs

As Erceg et al. reported (24), approximately half of the PhoRC peaks are located within promoter regions, while the remainder are in other genomic regions. Based on the physical overlap between PhoRC peaks with four different categories of genomic locations, the PhoRC peaks were categorized into four classes: promoters, enhancers, intergenic (excluding enhancers), and intragenic regions. *In vivo* assays of several overlapped PRE-enhancers reveal multifunctional PREs; for example, a PRE region can act as a (repressive) PRE in one tissue or developmental stage while the same region can act as enhancer at a different time and in different tissues (24). Thus, we use the same dataset and classification as described above to train a second, classifying RF model, in order to identify the key features distinguishing different PRE subtypes.

Based upon the approach of Erceg and colleagues, we classified the PhoRC binding sites in our training set into 4 categories. In order to have a balanced positive dataset of each of the potential PRE classes, we randomly selected 140 PhoRC binding locations (equal to the size of the minimum positive data set, *i.e*. intragenic) and tried to fit an RF model based on the motifs in Group A-D to predict the confidence score of each of these categories for each predicted potential PRE (Fig 6 and S10 Fig). The model performance on the withheld testing set, assessed using the AUC values under both ROC and P-R curves, demonstrates that among these four models, the model using Group A motifs has the best performance in categorizing the potential PRE regions, while Groups B and D show poor performance (S10 Fig). It is notable here that among the feature subsets in Groups B-D (all of which contain only some of the features in Group A), the strongest performance arises from Group C (Fig. S10 B&B’). Furthermore, the performance of this model, which completely excludes Polycomb-related motifs (Group C; Fig. S10 B&B’) is similar to or better than that including only Polycomb-related motifs (Group B; Fig. S10 A&A’), suggesting substantial importance for accessory proteins in defining the functionalities of different PREs. The performances of the Group A model on each classification group are compared in the confusion matrix plotted in Fig 7. We find that the model is essentially unable to identify intergenic regions, which may lack any uniquely defining characteristics within the feature set used here. Fig 8 and S11 indicates top 30 important features that are involved in categorizing the predicted PRE regions using models built on feature Groups A-D. These findings highlight the importance of accessory DNA binding proteins in determining the functionality of PREs, as the most informative features for distinguishing between different PRE subclasses tend not to be PcG and TrxG proteins. In order to obtain more insight on what variables are potentially important to distinguish four classified PRE/TRE regions, we looked at the rz-score distributions of 15 top variables and compare them with the distribution of target DNA binding association motifs (using the intergenic region as a base line) via a two-tailed Wilcoxon test (as in the case we simply seek distributions different from the intergenic region, rather than singling out *a priori* only cases that are enriched relative to the intergenic region). After deriving the p-values and applying FDR correction, we considered q-value less than 0.05 to indicate a significant difference between a given class and the intergenic region (Fig 9). Comparing these factors among the 4 different PRE classes shows that some of these variables like BEAF-32 and pad are informative of promoters, IPRM1 and IPRM2 (Cg and Trl based on Tomtom motif matching) in enhancers have significantly different average densities than other classes. Thus, the frequency of these protein binding sites may play an important role in establishing the functional identities of different regions (for example, in distinguishing enhancer sites from other intergenic PREs). In general, however, the patterns identified in our classification model defy simple interpretation (as is often the case for RF models).

**Figure 6.**
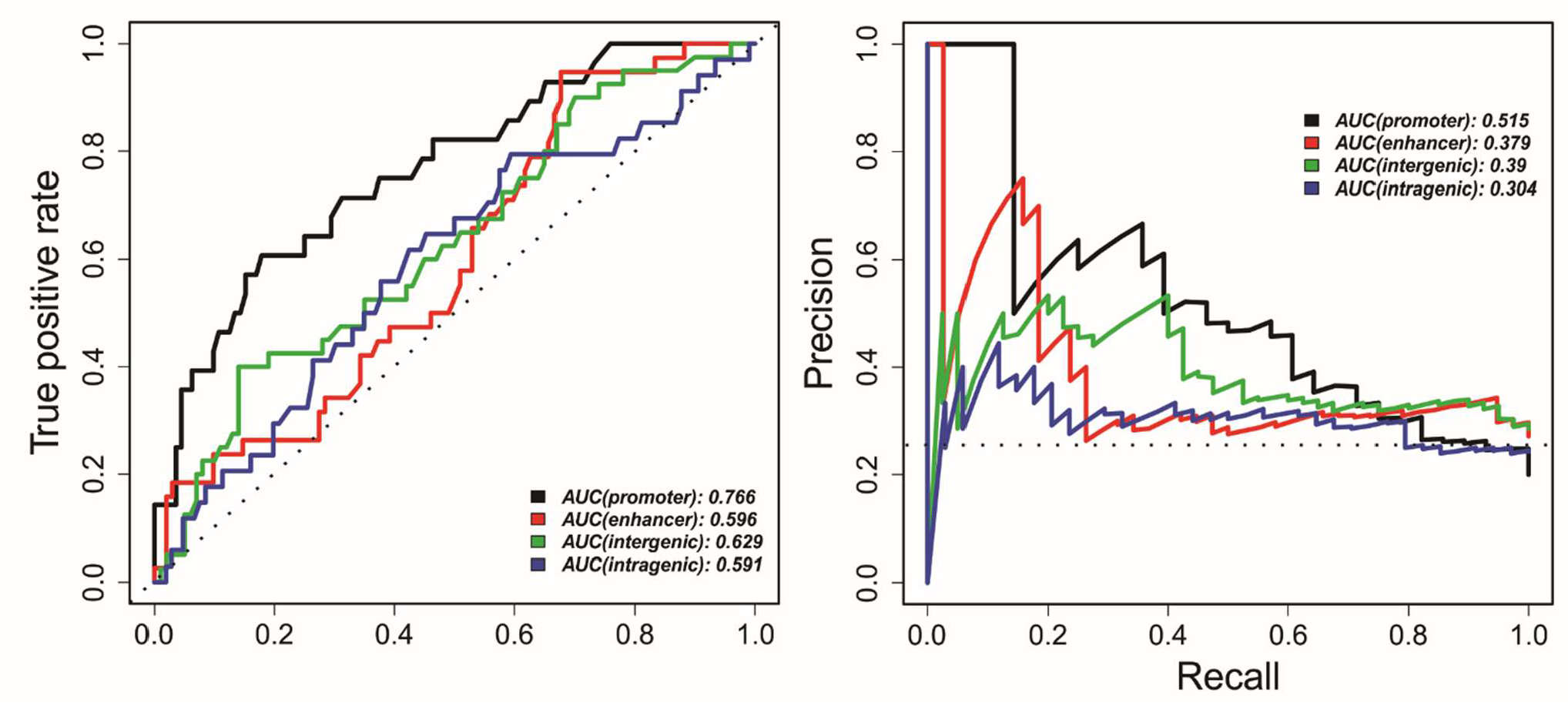
Evaluation of the performance of an RF model (based on Group A features) on the withheld testing data set. We assess the ability to categorize potential PRE regions using ROC (left) and P-R (right) curves. AUC indicates the area under each curve. Dashed lines indicate a hypothetical null model’s performance.

**Figure 7.**
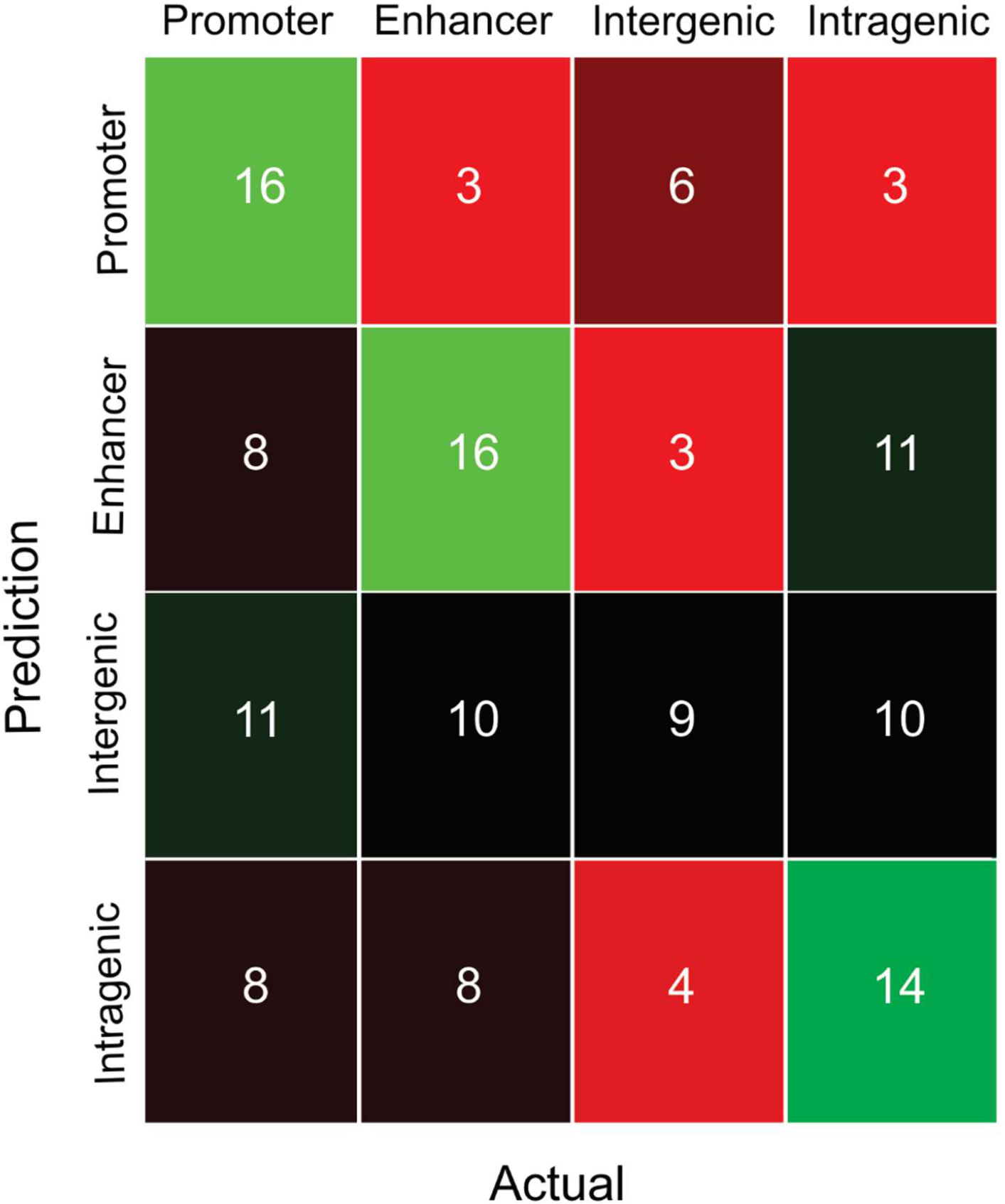
The heatmap of the confusion matrix for performance of 4 different models including promoter, enhancer, intergenic and intragenic (from Group A in Fig 6), showing the correlation between model predictions and ground-truth values for members of different classes in testing data.

**Figure 8.**
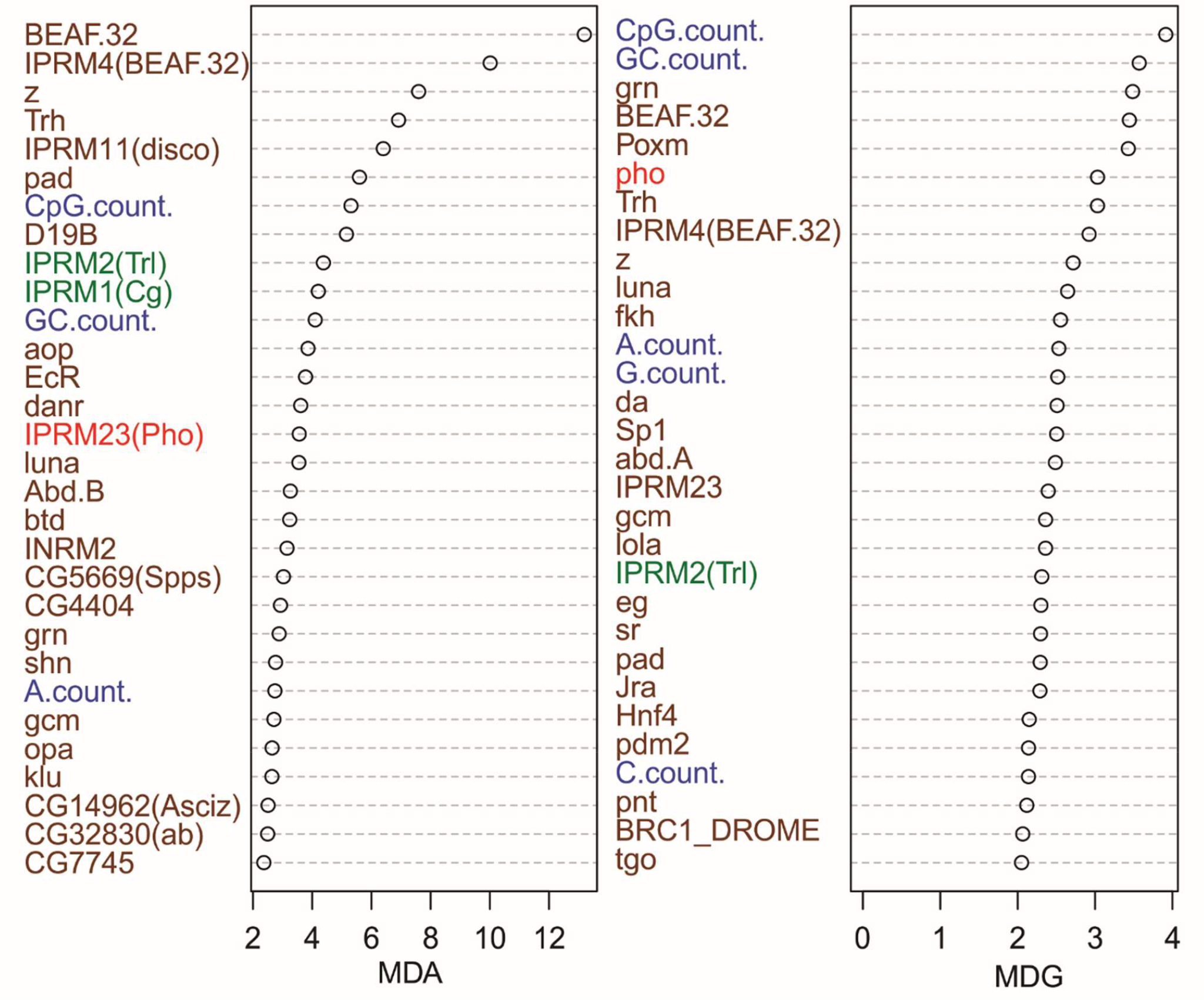
Variable importance table for the RF model to classify potential PREs which build based on all motifs. Mean decrease in accuracy (MDA) and mean decrease in Gini coefficient (MDG) of effective factors in RF model. Trl and Trl related motifs are shown in green, sequence features in blue, Pho and Pho related motifs in red, and other features (predominantly other motifs) in brown.

**Figure 9.**
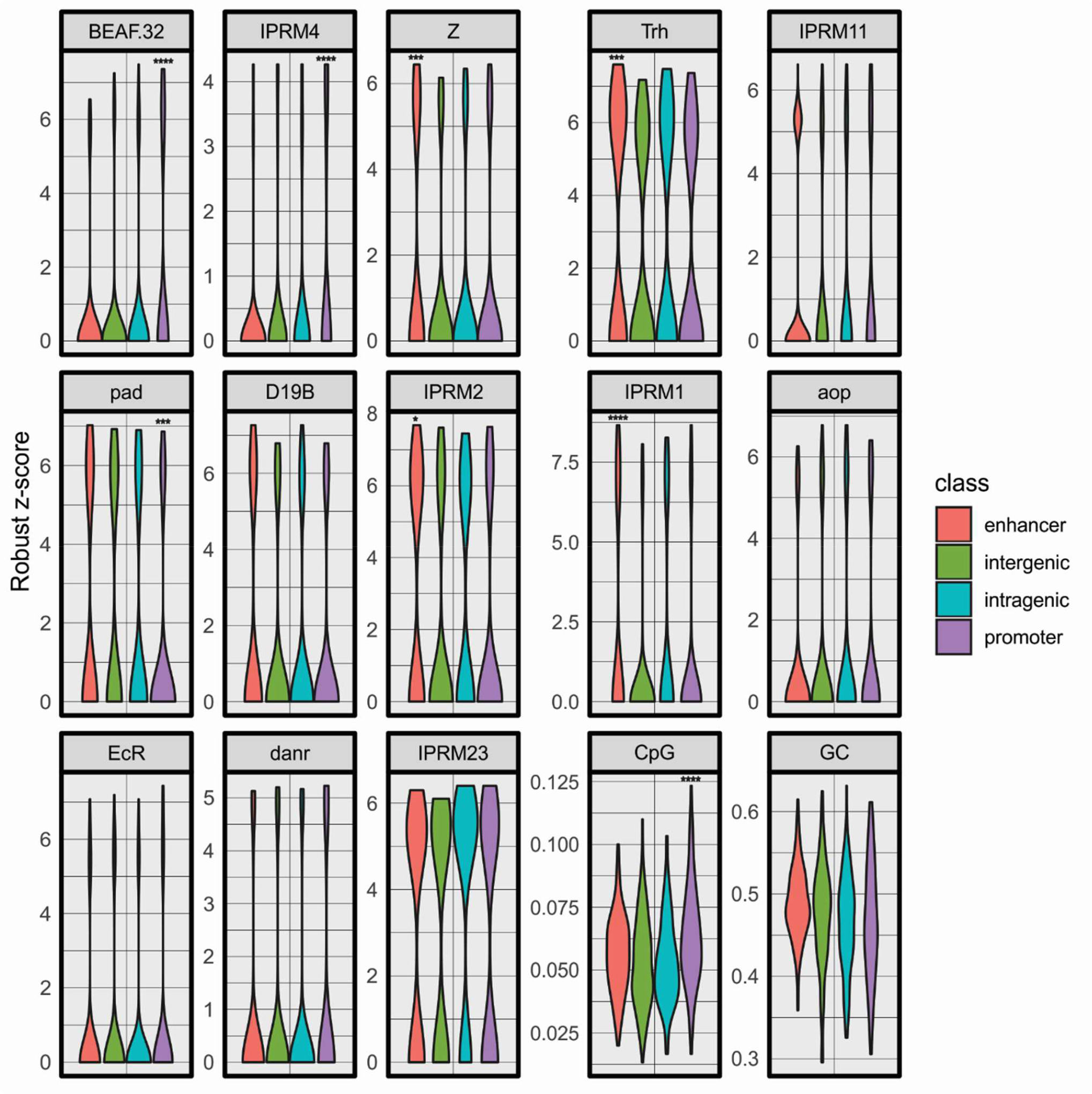
Distribution of rz-score of top 15 important variables in Fig 8 at different PhoRC binding classes. The two tailed Wilcoxon rank sum test q-values used to determine whether the distributions for promoter, enhancer, and intragenic sites are significantly different from intergenic regions (each tested separately). The stars are only show the levels of significance. If a q-value is less than 0.05 it is one star (*), less than 0.01 it is two stars (**), less than 0.001 it is three stars (***) and less than 0.0001 it is four stars (****).

Fig 10 provides an example view of the outputs of the two-stage optimized RF model developed in the present study. This figure represents a successful prediction of potential PRE regions and their corresponding classes/functions around the *bx* PRE located on chromosome 3R. The known bx PRE has overlap with enhancer regions(24). As shown in Fig. 10, our model successfully predicts a PRE overlapping the bx enhancer (confidence score 0.975 in the binary PRE prediction model), and that potential PRE is classified as an enhancer by our second-stage model (confidence score 0.361 in the four-way PRE classification model). Classifications of all predicted potential PREs from our model are given in S8 Table.

**Figure 10.**
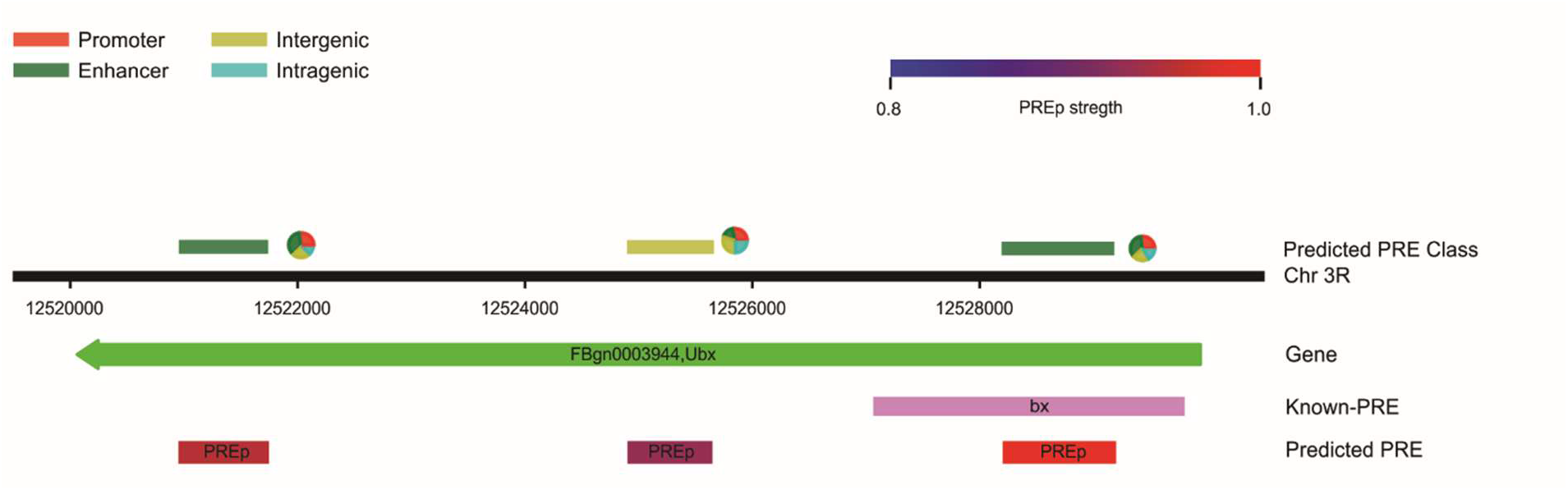
Example view of potential PRE prediction and their predicted category over entire *D. melanogaster* genome. The pie chart next to each prediction shows the fraction of trees as a confidence score for classifying target predicted potential PRE segment (cutoff greater than or equal to 0.8). The range of color from blue to red used to indicate the confidence of potential PRE (PREp) prediction in our binary model distinguishing PRE from non-PRE sites.

## Discussion

Based on a recent study of PhoRC locations via ChIP-seq assay (24), a large dataset of potential PRE regions became available to build a predictive model that would provide more general insight into PRE locations than prior efforts. Thus, we developed an RF classifier, using a robust machine learning algorithm with a large training set to predict potential PRE regions and their functions as enhancers, promoters, or both. Our model provided high performance in identifying potential PREs from non-PREs in our training set and enabled the identification of hundreds of other potential PREs throughout the genome. The identities of the most informative TFs, those that already known to be at PRE regions, involved in construction of the model to determine potential PRE regions over entire *D. melanogaster* genome. Removing the key TFs previously known to be involved in establishing PREs, such as Pho, Trl related motifs and also PhoRC, revealed the importance of other TFs in establishing active PREs, rather than PREs being dictated solely by the binding of core Polycomb complex components; likewise, reasonably good predictive performance could be achieved even in the absence of the motifs for the known Polycomb complex components. The annotated functions of these new TFs identified by our system revealed that they are crucial for key developmental processes. In two previous studies of genome-wide PRE prediction (26, 27), regardless of the type of the method and algorithm used, predictions were only based on sequence features and seven known common motifs in PRE regions, whereas we extended our study to consider the binding of all possible DNA-associated proteins that might contribute to establishing active PREs. Our results suggest that additional proteins beyond the known Polycomb complex core components are necessary to specify the locations, activities, and functions of active PREs in a condition-specific manner.

Validation of our model using experimental data sets that were not involved in model construction showed significant enrichments of overlap between our predictions and the experimental data sets, particularly for PREs with high confidence scores (>0.8 & >0.9). Of particular note, according to cutoff 0.8 and 0.9, there are 89 and 34 genes, respectively, which are shared between our high-confidence predictions and both experimental validation data sets, representing a particularly strong class of likely PRE-regulated targets which appear to drive developmental processes (S9 Table).

In interpreting our findings, it is important to consider that all of the data used in our model are centered around Pho and dSfmbt occupancy, specifically in mesodermal embryonic cells in 4-6 h and 6-8 h after egg laying, corresponding to the conditions used to generate our training data sets (24). Thus, our predictions are almost certainly condition specific and can only be considered to represent potentially active PREs in the absence of more definitive experimental. As Erceg and colleagues found, there are 994 locations on the genome that covered by both Pho and dSfmbt ChIP-Seq peaks at stages from 4-8h of embryogenesis. Among them, 468 of them are considered as promoters, 225 of them as enhancers, 161 as intergenic, and 140 of them as intragenic. The authors then selected 16 peak regions to test the PRE or enhancer activities, finding that 9 have dual functions and were able to either activate or silence transcription in a spatio-temporal manner. Notably, all of other key accessory DNA-binding proteins that appear based on our model to contribute to active PREs are expressed at 80-100% of their maximal levels at the time and tissue of the Erceg experiments except for sr (with 5.2%), Mad (with 57%), and grn (with 55%), using data from the BDGP database(47–49). It is certainly possible that under different conditions (developmental stage or cell types), different groups of accessory DNA-binding proteins would be active in promoting PRE formation at other potential positions on the genome; the identification of such factors, their application in PRE prediction, and experimental testing of their roles in establishing PREs in a time- and tissue-specific manner, will be a fruitful topic for future work.

Recent data have shown that PREs are able to act as silencing elements or enhancers in different cellular environments (24). According to the overlap of potential PRE regions identified from their ChIP-seq assay with enhancers and promoters, each DNA segment was classified as an enhancer, promoter, intergenic, or intragenic region (the last being a catch-all for sites not falling into one of the other categories) (24). Thus, we developed a secondary model to categorize predicted potential PRE regions based on their possible function to use it to identify key sequence features of the four different categories of PREs. We evaluated the function of each potential PRE region predicted in our original classifier using a combination of all motif and sequence features present in our potential PRE regions. Our findings extend the idea of Erceg *et al.*, who suggested that a number of known PREs, including bx, have dual functionality as repressors and enhancers. Thus, we used our regression model to characterize important factors in each potential PRE class (Fig 10) with particularly good performance at identifying PREs in promoters and in potential enhancers. Feature importance analysis then showed that several known chromatin remodeling TFs including “CTCF”, “BEAF-32”, and “Trl”, are among the most important features for classifying PREs using our RF model. It has already been shown that the activation by enhancers and repression by the PcG proteins are involved in higher-order interactions, either by looping or forming discrete domains of interaction (50–52). Thus, we compared our prediction results with the looping data obtained from Ogiyama *et al.* (53). In this study they identified higher order 3D chromatin organization using Hi-C experiments as well as Polycomb domain loops during *Drosophila* embryogenesis and determined active and repressive chromatin loops. Comparing the overlap of their hand-selected loops with our predicted PRE regions, we obtained enrichments of 11.83 (p=9.9e-5) and 42.56 (p=9.9e-5, name test) using our predictions with cut-offs of 0.8 and 0.9, respectively; these findings illustrate that our predicted PREs are strongly enriched among Polycomb domain dependent loops. In addition, comparison with yet other experimental data sets likewise elevates confidence in our prediction results. Follmer et al. (54) analyzed PcG protein localization, including Psc and Ph, in mitotic chromosomes in *D. melanogaster* S2 cells. The overlap analysis of Psc and Ph ChIP-seq peaks with predicted potential PREs show 8.3 and 6.7 fold enrichment at cutoff 0.8, and 22.1 and 13.42 at cutoff 0.9 (in all cases p-value= 9.9e-5 using permutation test), respectively.

Thus, taken together, we have been able to identify many new potential PREs (summarized in S8 Table using a cutoff confidence score of 0.8), and identified key features contributing both to the establishment of active PREs in mesoderm cell types for embryos 4-8h after fertilization, and features particularly useful in differentiating between PREs that might be active as enhancers, promoters, or involved in more general functions. One particularly important insight to be gained from this work is the identification of potential accessory DNA-associated proteins that might contribute to PRE formation, at least inasmuch as the presence of motifs for those proteins is strongly informative of PRE locations. Future experiments will be required to confirm what causal roles, if any, those factors play in actually establishing active PREs, and whether (as we hypothesize) different sets of factors play similar roles in establishing PREs in different tissues and developmental timepoints. In addition, future predictions and experiments should enable us to obtain similarly informative models of active PREs under different physiological conditions, ultimately assisting in the dissection of what factors trigger the changing locations and activities of PREs over the course of development.

## Methods

### Motif Inference and Scoring

We used meme-chip (55) to construct PWMs from ChIP-seq data and fimo (56) to calculate all possible biding scores of each PWM (x_i_) in increments of one base pair over *D. melanogaster* genome. *De novo* motif discovery was performed on the feature-separated *D. melanogaster* genome, version 5.9 obtained from flybase, 250bp +/− around the Pho/dSfmbt peak summits from the Erceg 2017 dataset using MEME-ChIP 4.11.1 (55), with the following parameters: “meme-chip -o meme-chip -meme-minw 10 -db PWM.txt -bfile bg2.txt ChIP-seq.fasta” while bg2.txt is A/T/C/G with equal frequency of 0.25, and PWM.txt is a database of all available DNA motifs, respectively. The motif discovery for nejire and psq ChIP-seq data, obtained from modENCODE (32, 33), was done using the same parameters except that the nucleotide frequencies were considered as 0.285 for each of A and T and 0.215 for each of C and G. To predict DNA-associated protein binding sites over the entire genome, we used the “find individual motif occurrences” (fimo) tool from MEME suite v. 4.11.1. To calculate the MAD and rz-score and calculate the distribution of the matching scores for each motif at each position, we used the following command, “fimo --oc fimo --max-strand --no-qvalue --bgfile bg.txt -- thresh 1 --max-stored-scores 10000000 --text --parse-genomic-coord pwm.meme dmelv5.9.fasta” where bg.tx contains frequencies equivalent to those used in nejire and psq motif analysis. We then calculated the position-wise rz-scores using the median and MAD, which is robust to single outliers, of a given population of fimo score across whole genome according to Equation 1. Only rz-scores with a value larger than 3 were retained.

### Construction of the training set

RF is a supervised learning algorithm. Thus, we needed to provide it with a training set containing both positive (PhoRC binding sites) and negative (non-PRE) examples. We constructed a sequence collection of PhoRC ChIP-seq data (24) as known PRE sequences, and the same number of random sequences were selected over the whole genome as a negative dataset. A few potential PRE locations of our constructed dataset (less than 1%) showed zero DNA-associated protein binding sites, thus we deleted any such sequences from our training dataset. The positive and negative examples were then split to 75% of the data for training the models and 25% of as testing dataset for models’ validation. The PhoRC peaks in the training ChIP-seq data were varied from 100 to ~1500 bps. Thus, we select ±150 bps around the summit of each peak in order to have all sequences have the same length. To ensure that our control sequences were randomly selected, we repeated the process of random sequence selection 10 times. For each selection we calculated the AUC of ROC and P-R curves (Fig S6) which were all within ± 0.03 of main RF model based on Group A classes. In order to predict potential PREs we scanned *D. melanogaster* genome v5.9 including chromosomes 3R, 3L, 2R, 2L, 4, U, Uextra, X both for both euchromatin and heterochromatin obtained from flybase(31), with a sliding window of 300 bps that incremented with a constant step of 50 bps. For each potential PRE sequence, we calculated sequence features and the possible predicted DNA-associated binding sites and applied the RF model trained above. At each cut-off, we identified as potential PREs all regions above that confidence score, and then expanded each PRE to cover the immediately flanking regions until the confidence score dropped to 0.5

### Random Forest parameters

Random forest is a nonparametric method that can also be used as a binary classifier. We used the RF as a classifier to predict whether a given locus is actually a potential PRE site. All calculations done here used the RandomForest R package (37). The RF model tuned via “TunRF” with the following parameters; ntreeTry=200, stepFactor=1.5, and improve=0.01. For final model, the parameters mtry = 17 and ntree = 200 were used to fit the model. The following parameters and command were used to create RF model:

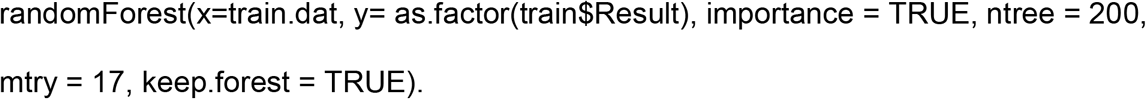

### Model Validation

We used a fivefold cross validation to validate our prediction model. The total training data set was divided into 5 equal size subsets with randomly selected values without replacement. At each time, 4 of the subset were used as a training set, whereas remaining subset formed the corresponding testing set. This procedure was repeated five times with every subset selected once for training. For further validation we also constructed 10 different datasets. The positive section of these datasets all were the same while the negative part of each dataset was randomly picked up over *D. melanogaster* genome. In each case, 75% of each dataset was used for training the model and 25% as testing sets. To avoid retraining the testing sets, we kept the positive part of all testing sets the same as the testing set that we used to validate our main model (based on Group A) (Fig S6, A&A’). Moreover, we applied the main model on each of these 10 different test sets and it showed the similar performance to each of these 10 different models (Fig S6, B&B’).

### Permutation test for gene set overlap

To evaluate the statistical significance of the overlaps between two or more gene lists, we implemented a simple permutation test. For each of 10,000 samples, we randomly picked different regions of the genome of the same number and length as our true PREs, as a null set of PRE-predictions. We then assigned regulated gene lists to the null samples following the same procedure as was used for the actual PRE-predictions and took the fraction of null samples with equal or greater overlap than they actually observed overlap as the p-value.

