## Supplemental Figures and Table Captions for "Genome-wide Prediction of Potential Polycomb Response Elements and their Functions"

**Table S1.** Candidates motifs identified from ChIP-seq data, obtained from modENCODE and analyzed by meme-chip. These motifs are not existing in CIS-BP database. This table provided as a separate file as txt format.

| Motif enrichment analysis ofChIP-seq data (Erceg et al.2017) | | |
|---|---|---|
| Arbitrary motif name | Sequence logo of the positionweight matrix | Motif name according to TOMTOM |
| PhoRC | 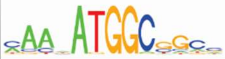 | Pho-Sfmbt complex |
| IPRM1 | 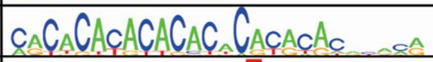 | Combgap |
| IPRM2 | 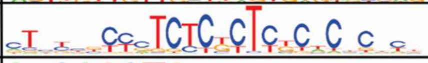 | Tomtom: Trl |
| IPRM3 | 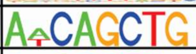 | Tomtom: nau |
| IPRM4 | 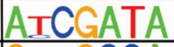 | Tomtom: BEAF32 |
| IPRM5 | 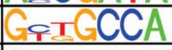 | Tomtom: FBgn0038766 |
| IPRM6 | 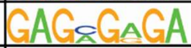 | Tomtom: Trl |
| IPRM7 | 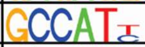 | Tomtom: Phol |
| IPRM8 | 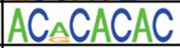 | Tomtom:dar1 |
| IPRM9 | 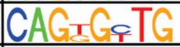 | Tomtom: Zif, Zipic |
| IPRM10 | 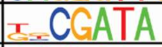 | Tomtom: srp , BEAF32 |
| IPRM11 | 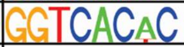 | Tomtom:disco |
| IPRM12 | 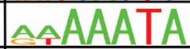 | Tomtom: Slp1, Jigr1 |
| IPRM13 | 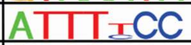 | Tomtom: Lola, dl |
| IPRM14 | 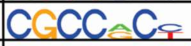 | Tomtom: SREBP, brk |
| IPRM15 | 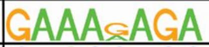 | Tomtom: Trl , Blimp1 |
| IPRM16 | 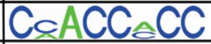 | Tomtom: ttk, klu |
| IPRM17 | 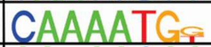 | Tomtom: Pho, Phol |
| IPRM18 | 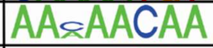 | Tomtom: Aef1 |
| IPRM19 | 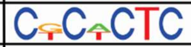 | Tomtom: Trl |
| IPRM20 | 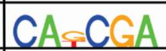 | Tomtom: Not Recognized |
| IPRM21 | 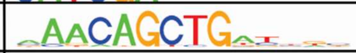 | Tomtom: crp |
| IPRM22 | 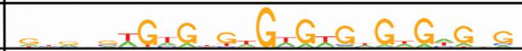 | Combgap |
| IPRM23 | 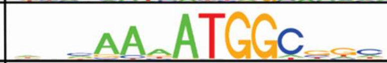 | Tomtom: Pho |
| IPRM24 | 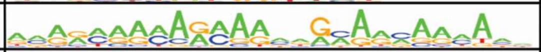 | Tomtom: rn, jim |
| IPRM25 | 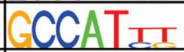 | Tomtom: Pho |
| IPRM26 | 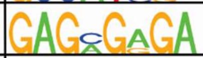 | Tomtom: Trl |
| IPRM27 | 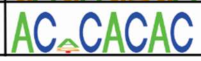 | Tomtom: dar1 |

**Figure S2.** Candidate motifs and likely identities of corresponding binding proteins, identified from ChIP-seq data set from Erceg et al. (1) via meme-chip. The PhoRC and Cg motifs are the same as in Erceg et al. (1) and Ray et al. (2). The rest of the motifs predicted via Tomtom(3) from the MEME Suite.

**Table S3.** Predicted transcription factor binding site database on chromosomes X, 4, 3R, 3L, 2R and 2L, U, Uextra and their related heterochromatin separately. The RF features were extracted from this database. Due to the size of the database, the files can be downloaded as a compressed archive from https://drive.google.com/file/d/1IXIeWKv0Jd-6YDj4fNPXOeALxNHOC2lg/view?usp=sharing

**Table S4.** A list of DNA-associated protein motifs that bind to PhoRC ChIP-seq peaks *within +/- 150 bps of peak summit. This table provided as a separate file as excel format.

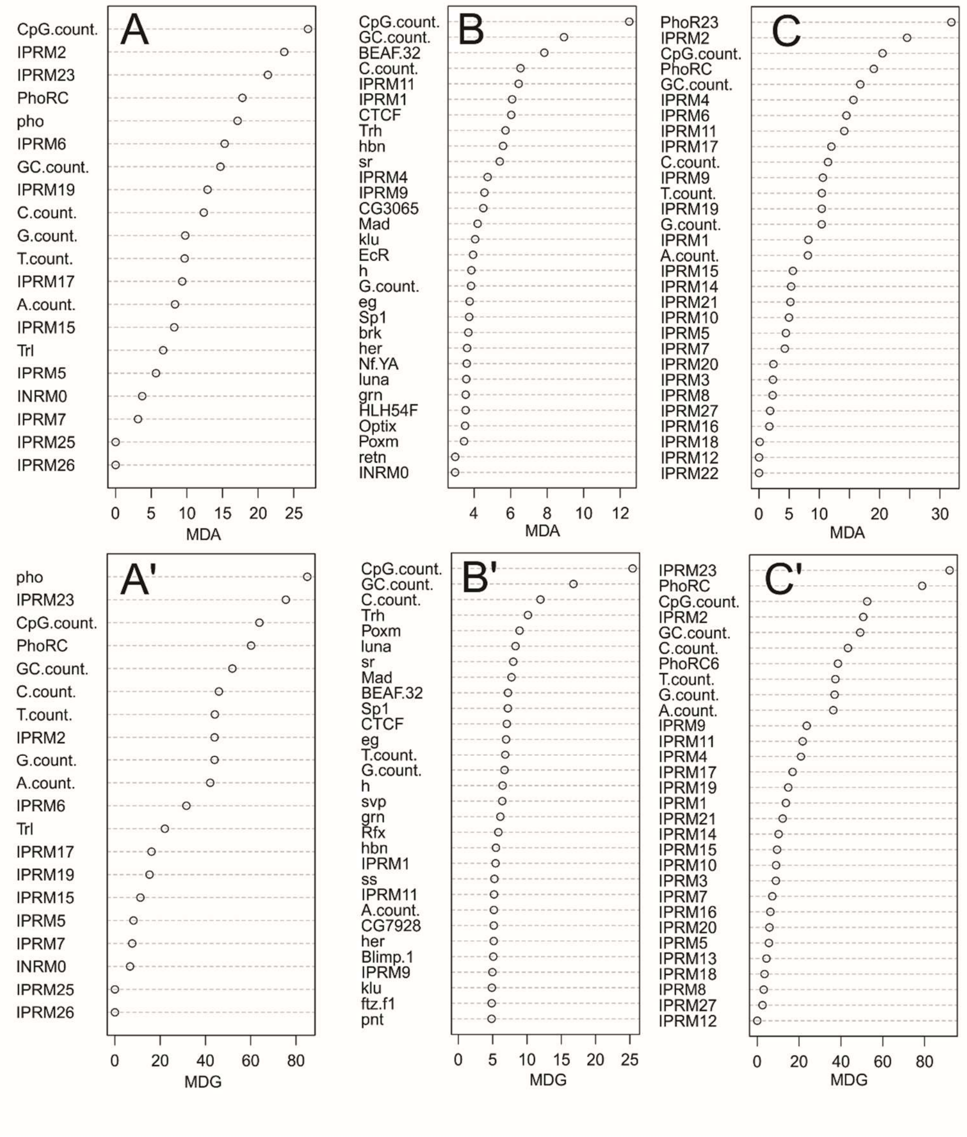


**Figure S5.** Extended variable importance plot from Fig 3. Top plots show the mean decrease in accuracy (MDA) and the bottom plots show mean decrease in Gini index (MDG). (A&A') Variable importance plots for Group B model based on Pho, Pho related motifs, Trl and Trl related motifs only as mentioned in Fig 3, (B&B') Group C model which considers every motif except motifs in Group A, (C&C') Group D model which built upon the motifs inferred from ChIP-seq assays as listed in S2 Fig. In all cases the elementary sequence properties (e.g., nucleotide composition) are also included as features.

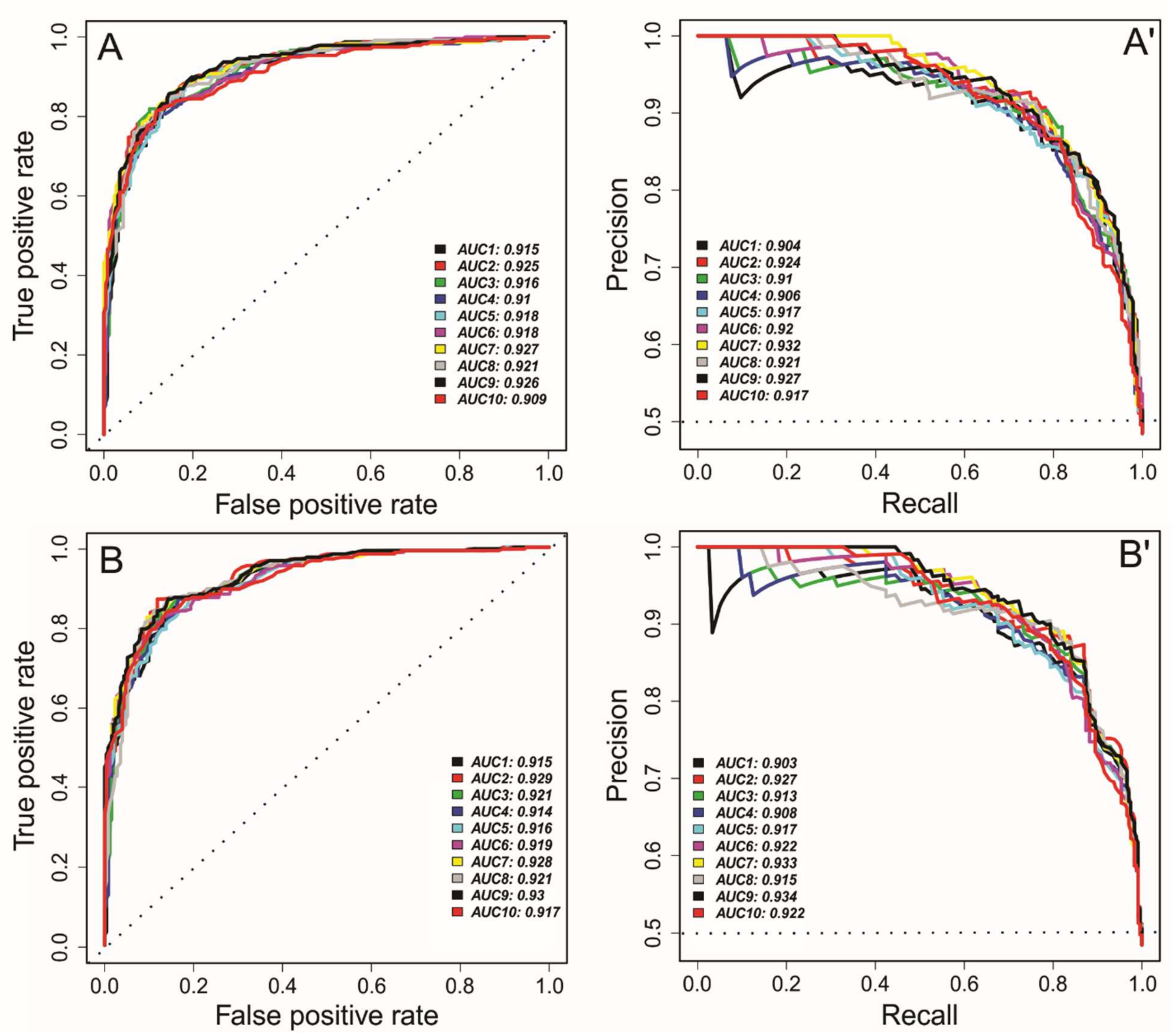


**Figure S6.** The top plots, ROC (Panel A) and P-R (panel A'), showing the performance of 10 models - with same positive and different negative data sets- based on the withheld testing sets. The bottom plots, ROC (Panel B) and P-R (panel B'), show the performance of the main model (the model built based on Group A and used throughout the main text) on the 10 different testing data sets used in panels A-A' without any further optimization.

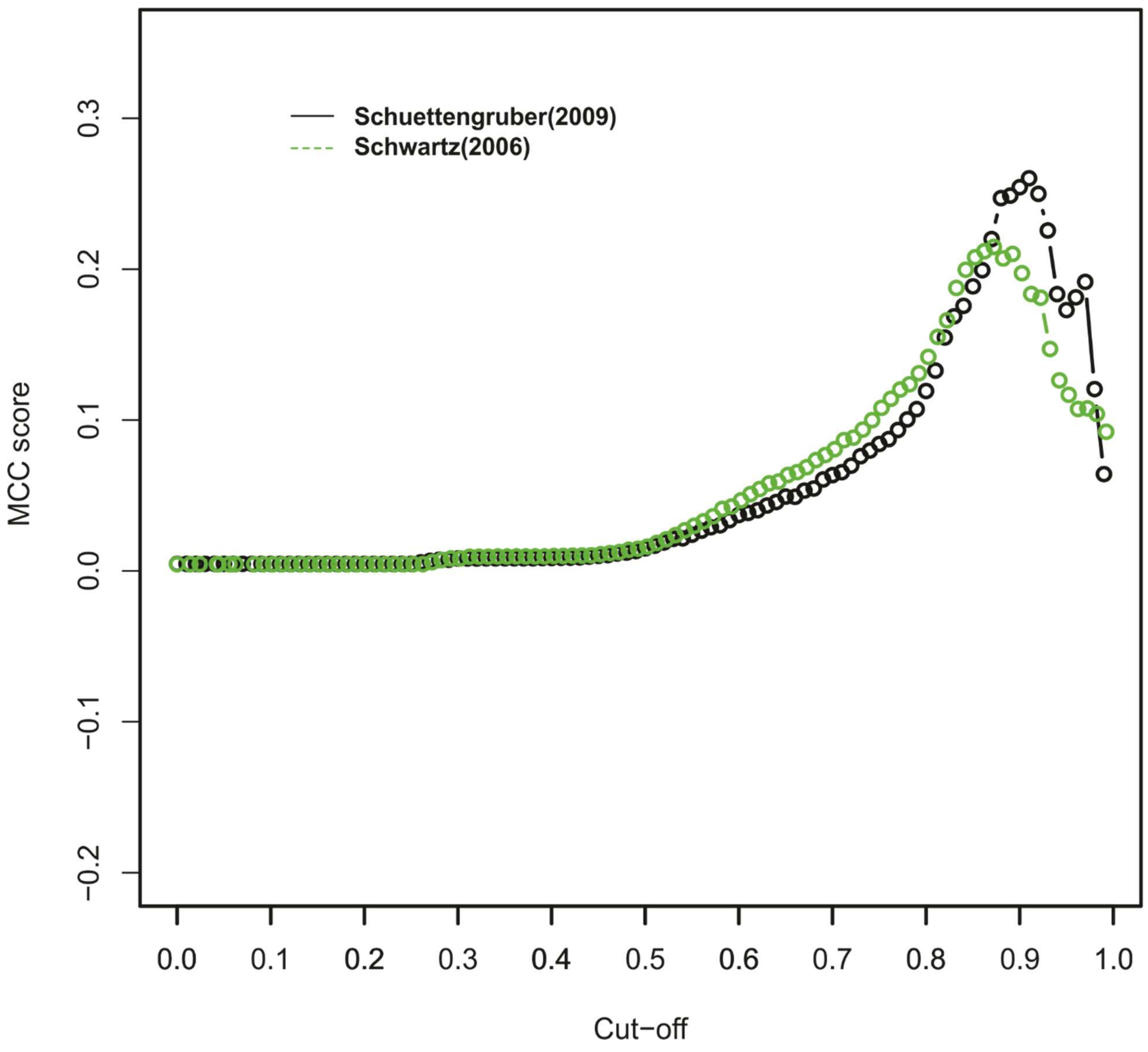


**Figure S7.** Optimization of random forest threshold based on overlaps of PRE calls with withheld experimental data sets. See main text for references. Plotted is the Matthews correlation coefficient (MCC;(4)) based on the overlap between the list of PRE target gene lists from Schuettengruber et al.(5) and Schwartz et al.(6) with the target genes of potential PRE regions in our model.

**Table S8.** Predicted potential PRE regions and their classes together with their corresponding confidence score (including all regions with a confidence score at the peak above 0.8). The file is in gff3 format. The last field contains both the confidence score of the prediction of a region as a PRE (PREp), and the confidence scores for each of the four PRE types from the classification model. This table provided as a separate file as gff3 format.

**-Table S9.** Listing of predicted PRE-regulated genes, as well as target genes from experimental studies, as described in the text accompanying Fig. 5. The “predicted genes at cutoff 0.8 and 0.9” tabs, show list of predicted PRE target genes based on our RF model at a confidence score cutoff of 0.8 and 0.9, respectively. The “34 and 89 overlapped genes cutoff=0.8 and 0.9” tabs include list of genes which are commons among Schuettengruber et al.(5) and Schwartz et al.(6) and our prediction genes (each of the experimental data sets are also shown in separate tabs). This table provided as a separate file as excel format

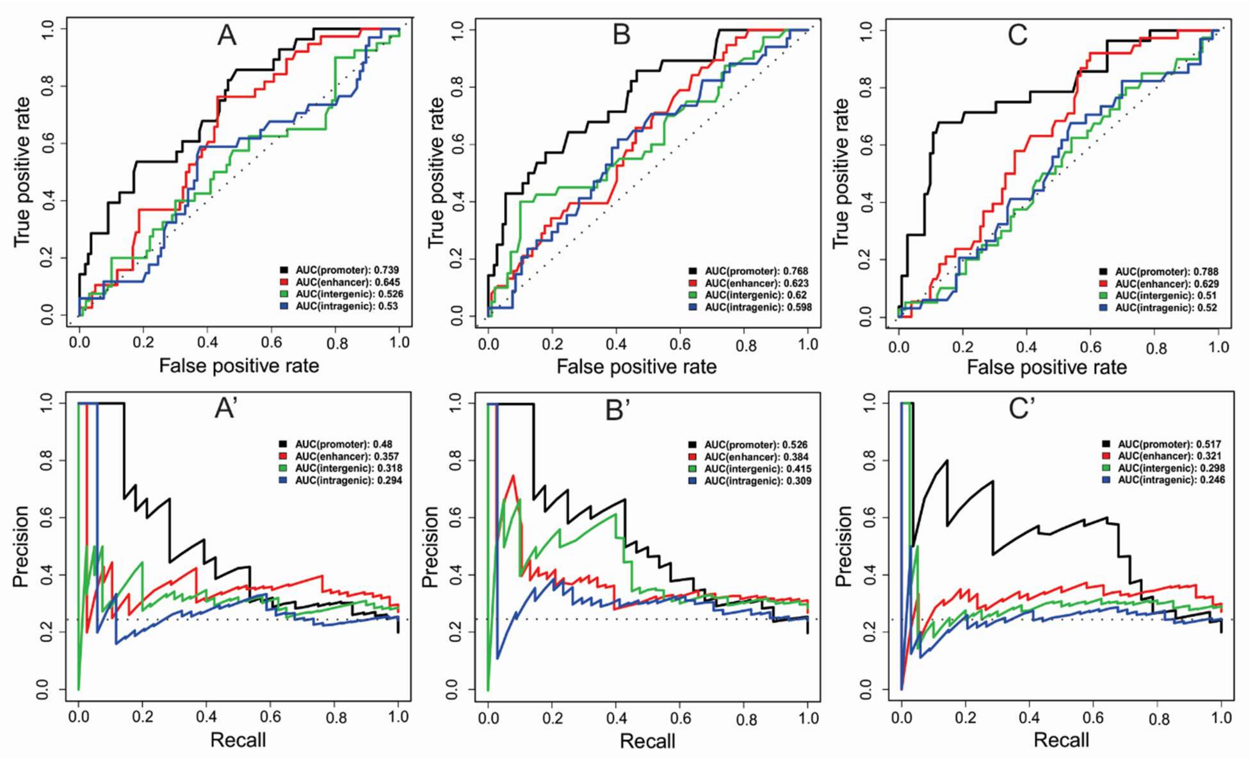


**Figure S10.** Extended ROC (top) and P-R (bottom) plots equivalent to Fig. 6, using models based on Group B (A&A'), Group C(B&B') and Group D(C&C') features. The top plots indicate the ROC while the bottom plots show P-R curves. (A&A') The performance plots based on the Group B features; (B&B') model based on Group C features; (C&C') model based on Group D features (see main text for group definitions).

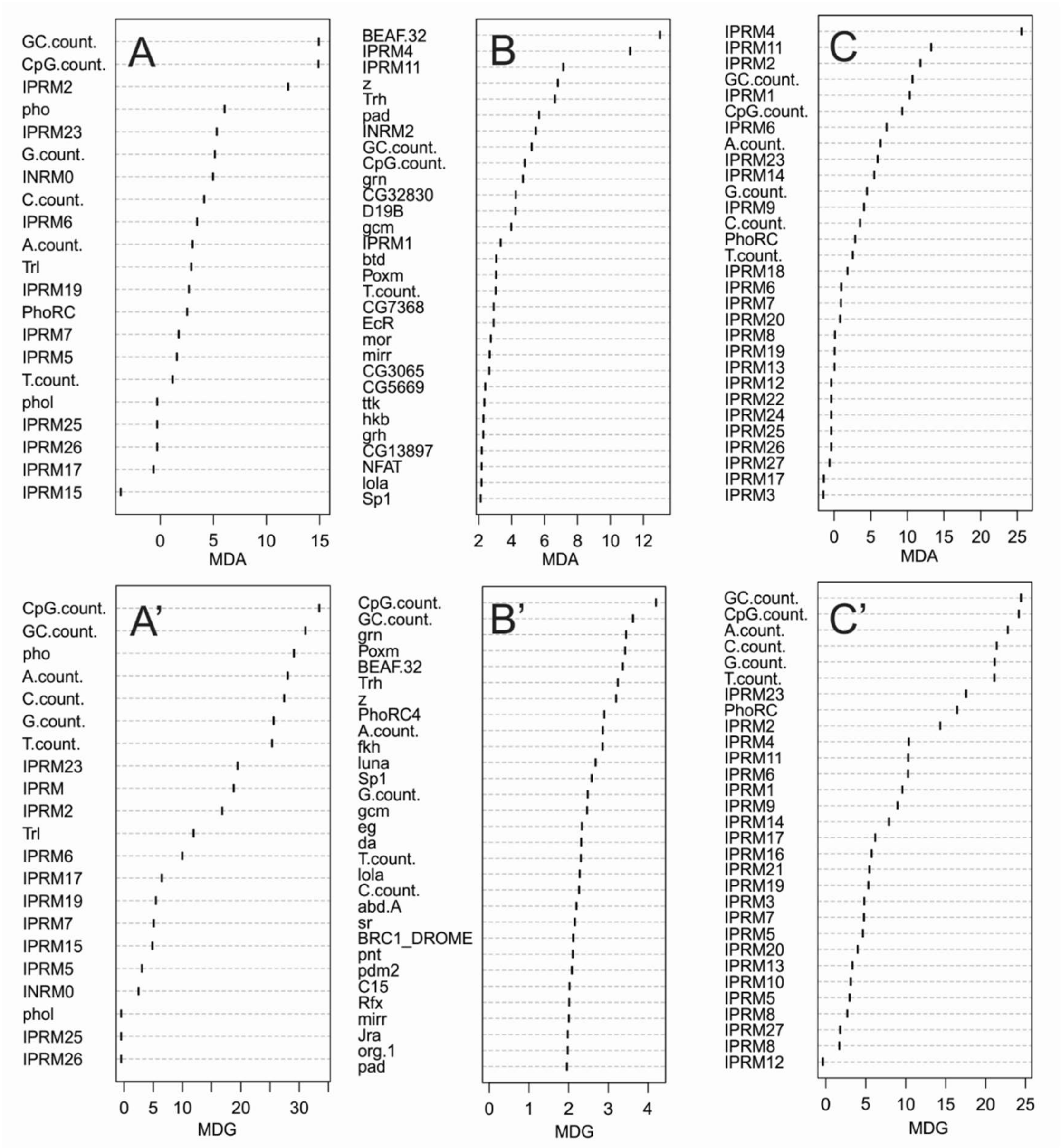


**Figure S11.** Extended plot from Fig 8. Mean decrease in accuracy (MDA) (Top) and mean decrease in Gini coefficient (MDG)(Bottom) of effective factors in RF model. Variable importance table for (A&A’) Group B model ;(B&B’) Group C model; (C&C’) Group D model.
